# Respiratory Phase Orchestrates Human Olfactory Cortical Dynamics

**DOI:** 10.1101/2025.09.25.678425

**Authors:** Alisa Y. Hathaway, Todd P. Coleman

## Abstract

Olfaction is fundamental to perception, memory, and emotion, yet the neural basis of human odor processing remains poorly understood. Here, we demonstrate that odor-evoked brain activity is organized by respiratory phase–not absolute time–revealing that natural breathing structures olfactory cortical dynamics. Using simultaneous nasal respiration and electroencephalography (EEG) recordings in 17 participants, we delivered naturalistic odors (peppermint, oregano, and grapefruit) synchronized to individual breathing cycles. Odor presentation elicited distinct, respiratory phase–locked event-related spectral perturbations (RPL-ERSPs) in frontal and temporal regions, emerging during the first breath after stimulus onset. The most discriminative neural signatures were theta-band activity during inhalation and alpha-band activity during exhalation, patterns that were obscured in traditional time-locked analyses due to natural variability in breath duration. Leave-one-subject-out cross-validation of RPL-ERSPs achieved reliable odor discrimination (peppermint: AUC = 0.78; oregano: AUC = 0.68; grapefruit: AUC = 0.61), establishing that phase-locked features generalize across individuals. To connect neural responses to perceptual experience without relying on precise odor naming—which is notoriously difficult and can be disproportionately impaired in some populations—we quantified odor recognition using free-response descriptions scored with a semantic-similarity approach. Behavioral recognition strength paralleled the neural decoding hierarchy across odors, establishing the perceptual validity of RPL-ERSPs. RPL-ERSPs also revealed rapid habituation, with decoding performance declining over repeated presentations, due to selective degradation of the most informative phase-locked features. Together, these findings establish respiratory rhythm as a temporal scaffold for human olfactory cortical processing, positioning RPL-ERSPs as a sensitive, task-free approach for assessing olfactory function. We provide an open-source, respiration-synchronized olfactometer design (<$500) to enable replication and broader adoption of this approach.

**Teaser:** Aligning brain activity to natural breathing unlocks objective neural markers of human smell perception.

## 2 Introduction

Olfaction plays a central role in perception, memory, and emotion, and its impairment is a hallmark of several neurological and psychiatric disorders, including Parkinson’s, Alzheimer’s disease, eating disorders, and post traumatic stress disorder [1–4] – often appearing years before other symptoms. However, we lack sensitive paradigms for measuring olfactory deficit that can be deployed across diverse populations. Instead, it is common to use forced-choice paradigms, drawing from a fixed, perhaps culturally-biased, repertoire of scents (e.g. University of Pennsylvania Smell Identification Test, UPSIT [5]). Recent olfaction research has leveraged electrophysiology to uncover generalizable neuro-markers of odor processing, but feature experimental paradigms with rigid, volitional breathing instructions (e.g., timed inhalations, breath-holding [6, 7]) which may be challenging to consistently deploy. Moreover, forced inhalation may not use the same odor processing pathways as naturalistic, spontaneous odor perception, thus obscuring (potentially) more ecologically valid indicators.

The relevance of naturalistic respiration is underscored by a growing body of evidence showing how nasal breathing actively shapes brain activity [8]. For instance, respiration-synchronized oscillations have been measured in the rat olfactory bulb, piriform cortex, hippocampus, prefrontal cortex, and primary sensory areas like the barrel cortex [9, 10]. In humans, an intracranial electroencephalography (iEEG) study found that naturalistic nasal breathing synchronized slow (0.16–0.33 Hz) oscillations in the piriform cortex, amygdala, and hippocampus, and that memory retrieval performance was phase-dependent[11]. These rhythms are hypothesized to act as a global brain clock, coordinating activity across distant regions [12]. In olfaction research, a recent iEEG study measured elevated coupling between sniff phase and theta band amplitude in the piriform cortex during odor presentation [13]. However, forced inhalation paradigms may disrupt these phenomena, yielding a misleading snapshot of brain activity during olfactory processing.

In order to bridge this methodological gap, we designed a closed-loop olfactometer system that synchronizes scent delivery with naturalistic respiration, while reducing the cost from $30,000 [14] to <$500. This system delivered odorants in real time synchronized to the individual’s nasal respiration phase, without disrupting natural sniff patterns. When paired with electroencephalography (EEG), the system allowed us to probe central questions about respiratory-olfactory integration:

- Do odor-induced neural responses — such as beta, alpha, or theta activity — vary systematically depending on the phase of the respiratory cycle?
- Can respiration phase-locked neural analyses reveal dynamics that are obscured in traditional time-locked approaches?

However, respiration-synchronized nchronized protocol described below, wneural measurement addresses only one side of the problem: interpreting neural dynamics also requires behavioral measures that sensitively reflect what participants perceived. In olfaction, this is nontrivial because “identification accuracy” can reflect not only sensory evidence but also naming, familiarity, and semantic access [15, 16]. A parallel challenge in olfactory research concerns how perception is quantified behaviorally. Standardized identification tests (e.g., forced-choice paradigms used in clinical screening) yield convenient binary scoring [5]. However, they often conflate perceptual sensitivity with lexical access and familiarity, and can therefore underestimate odor knowledge in populations where labels are uncertain or culturally variable. In natural settings, odor knowledge is frequently expressed through flexible, context-driven descriptors, which can reflect meaningful recognition even when the canonical “label” is not produced [16, 17]. Accordingly, in addition to neural measures, we examined whether participants’ qualitative odor descriptions covaried with neural discriminability.

After the experiment concluded, participants provided free-form verbal descriptions of each odor stimulus. To quantify these open-ended responses without reducing them to correct/incorrect labels, we implemented a hybrid scoring approach combining professional odor descriptors from an empirical database with semantic similarity derived from modern language models [18]. This framework assigns partial credit to semantically appropriate but non-canonical descriptors, enabling a graded behavioral metric that is more aligned with how odors are described outside the laboratory. Such continuous scoring may be particularly valuable for aging populations and cross-cultural contexts, where exact label knowledge can vary even when perceptual recognition remains intact [19, 20]. Finally, treating behavior as a graded variable enables a stronger convergent test: if respiration-locked neural signatures truly reflect perceptual processing, they should scale with behavioral odor knowledge rather than only binary accuracy.

Using this system with 17 healthy participants exposed to naturalistic odors (peppermint, oregano, and grapefruit), we investigated whether neural oscillatory responses to odors were organized by respiratory phase rather than absolute time. We hypothesized that aligning neural data to respiratory phase would reveal structured, phase-locked dynamics that generalized across trials and individuals – dynamics that would be obscured or variable when analyzed relative to stimulus onset alone.

## 3 Methods

### 3.1 Participant Screening and Eligibility

Seventeen healthy volunteers (7 male, 10 female; ages 24-26) participated after providing informed consent under a Stanford University IRB protocol. Two participants scoring below mild microsmia on the University of Pennsylvania Smell Identification Test (UPSIT) were excluded to ensure normal olfactory function. Data were collected using a 128 Channel Geodesic EEG System at Stanford University’s Koret Human Neurosciences Community Laboratory. All subjects were asked to refrain from consuming caffeinated beverages for at least 2 hours before the experiment. They were asked not to wear perfume/cologne/other scented products on the day of the experiment[21].

### 3.2 EEG Data Collection

During the experiment, subjects were asked to sit still with their eyes open fixated on a central cross and to breathe normally through their nose. White noise played continuously through noise canceling earbuds to eliminate auditory confounds.

This passive experiment consisted of a no-odor (deionized water) control alongside three distinct odor stimuli: peppermint, oregano, and grapefruit. These odors were selected for their pronounced chemical differences. Specifically, grapefruit’s characteristic scent is primarily attributed to limonene, peppermint’s to menthol, and oregano’s to carvacrol [22–24]. While both limonene and menthol belong to the terpene chemical family, menthol is uniquely distinguished from typical terpenes by its ability to activate the trigeminal nerve, eliciting a cooling sensation [25], whereas limonene produces a characteristic fresh, citrus sensation [26]. Carvacrol, in contrast, is a member of the phenol chemical family, and elicits a warm/spicy sensation [27]. A summary of these chemical and sensory properties is provided in Table 1.

**Table 1:** Odor stimuli characteristics and chemical properties.

| Odor | Main Chemical | Chemical Family | Sensation | Trigeminal |
| --- | --- | --- | --- | --- |
| Peppermint | Menthol | Terpene | Cooling | Yes |
| Oregano | Carvacrol | Phenol | Warm/Spicy | No |
| Grapefruit | Limonene | Terpene | Fresh/Citrus | No |

For odor presentation, 2 mL of the specified doTERRA essential oil odors [28] were added to individual odor containers, each containing 100 mL of deionized water. Odors (or the no-odor control) were delivered using the respiration-synchronized protocol described below, with a randomized 15 s–30 s inter-stimulus interval and control-air flushing between trials to minimize carryover and reduce predictability. This procedure ran continuously for 30 min. The raw trial count, prior to cleaning and preprocessing, ranged from 20 to 30 per stimulus. This passive experiment was designed to not rely on active instructions or actions. After the experiment concluded, the experimenter asked the subjects to describe what they smelled, if anything. These free-form responses were later used to assess identification accuracy of participant responses for each odor.

### 3.3 Olfactometer Design and Operation

To enable respiration-synchronized odor delivery during natural, spontaneous nasal breathing, we developed a system built around a commercial air compressor nebulizer (Fig.1 **A**) [29], providing controlled airflow at 3 L/min to prevent nasal discomfort [30, 31]. The system delivered three distinct odorants plus a water control using glass bubble humidifiers constructed from Ball^®^ mason jars filled with essential oils diluted in deionized water (DI water; Fig.1 **D**). Air flowed through individual branches regulated by flowmeters (0-5 L/min; Fig.1 **C**), then through 3-way solenoid valves that directed unused scented air outside the experimental room to prevent ambient contamination (Fig.1 **E**). Final delivery occurred via a nasal cannula (Fig.1 **G**), which allowed precise bilateral nostril delivery while minimizing respiratory artifacts associated with cued or paced inhalations [32, 33].

#### Electronics and Control

An Arduino Uno microcontroller (Fig.1 **F**) orchestrated valve timing and communicated with the EEG system via electronic synchronization signals (TTL pulses) for precise event marking (latency ∼3.4 $s [34]). Solenoids were battery-powered to eliminate 60 Hz electrical noise that contaminates EEG recordings. Protective diodes prevented back-current damage during valve deactivation.

#### Respiration Monitoring

Real-time respiratory phase was detected with a Bosch BME280 temperature/pressure/humidity sensor embedded in the nasal cannula between nostrils (Fig.1 **H**). Because nasal airflow changes precede chest movement [6, 11], this placement provides superior temporal precision for olfaction studies. Respiratory phase transitions were identified by calculating derivatives within a rolling window of humidity measurements. Inhalation onset was detected when derivatives transitioned from positive to negative at the humidity peak, followed by sustained negative derivatives. Exhalation onset was identified when sustained positive derivatives indicated increasing humidity from exhaled air. Respiratory data were saved to a computer (Fig.1 **B**). An example respiratory response for one subject and one trial approximated by the nasal humidity measurements, is shown in Fig. 1, along with the synchronized TTL pulse, displayed as the dashed line.

**Figure 1.**
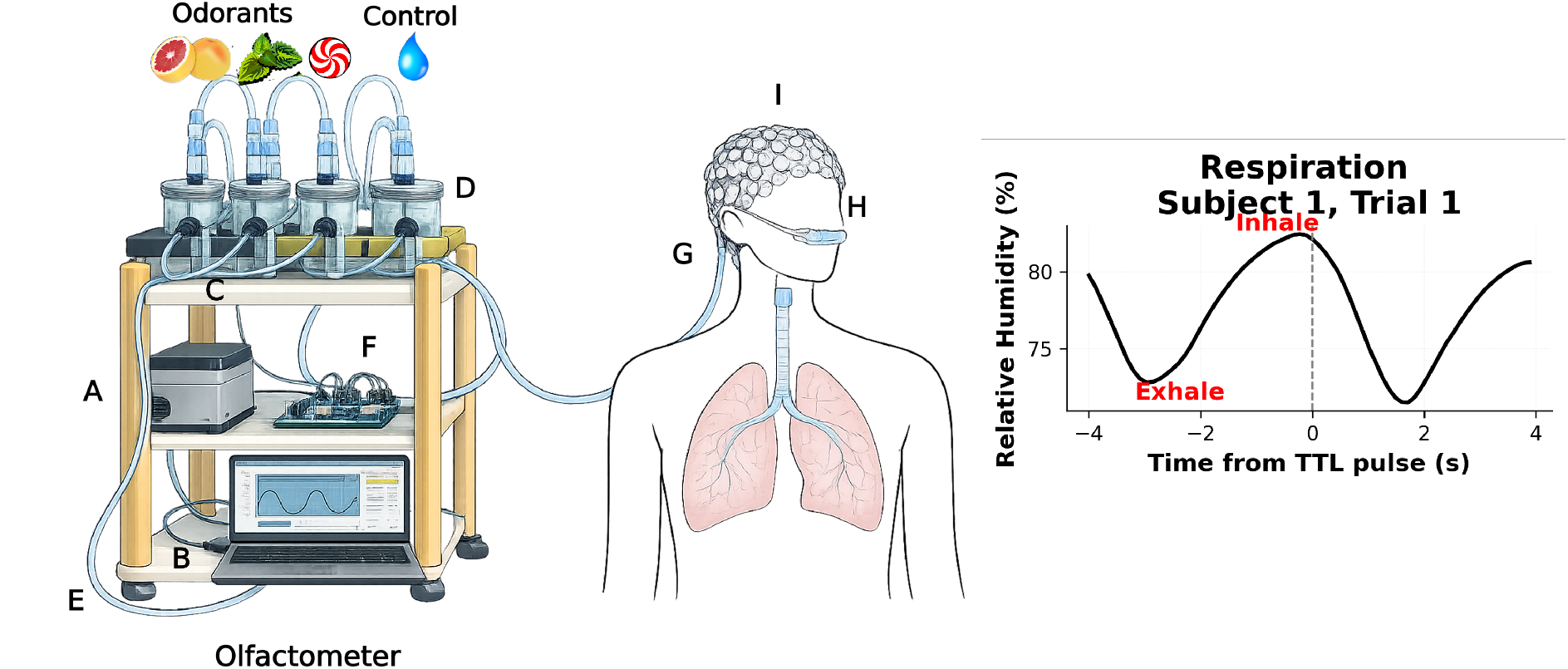
Experimental Setup and Hypothesized Respiratory Phase-Locked Neural Dynamics. **A:** Air compressor; **B:** Serial communication interface; **C:** Flow meters; **D:** Odorant containers; **E:** Tubing for excess air; **F:** Control circuitry; **G:** Nasal cannula; **H:** Respiratory Flow Sensor; **I:** Electroencephalography (EEG) cap. The right panel illustrates an example respiratory response for one subject and one trial approximated by the nasal humidity measurements. The synchronized TTL pulse is displayed as the dashed line, and inhale (peak) and exhale (trough) are labeled in red.

**Figure 2.**
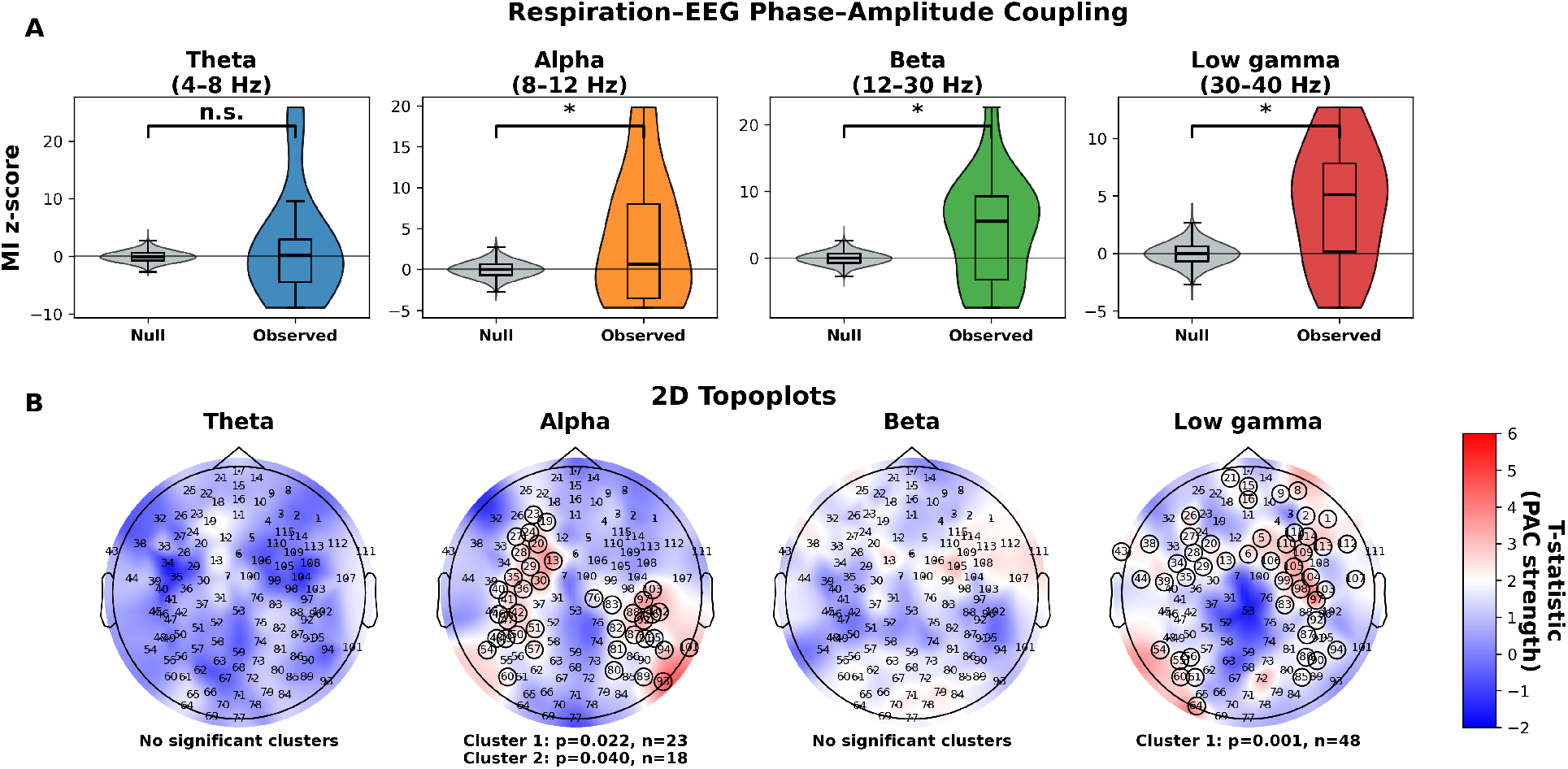
Respiratory phase modulates cortical oscillations with regional and frequency-specific patterns. **A:** Phase-amplitude coupling (PAC) strength between respiratory phase and EEG amplitude across frequency bands. Violin plots show z-scored modulation index (MI) distributions for null permutations (grey) and observed data, averaged for the whole head. Significant coupling (p<0.05), observed in alpha, beta, and low gamma bands. **B:** 2D topographic maps of the t-statistic with significant cluster electrodes circled.

#### Synchronized Delivery Protocol

The system operated in four stages: (1) continuous breath detection via temperature/humidity monitoring, (2) odor preparation following a randomized 15 s-30 s inter-stimulus interval by activating the first solenoid for up to 10 s, (3) timed release upon detection of exhalation, which allowed scented air to traverse the 1.21 m cannula and arrive at the nose at the onset of the subsequent inhalation, and (4) TTL pulse transmission to the EEG system marking the precise timestamp of inhalation onset (scent arrival). This design ensured odors reached participants at the optimal respiratory phase without disrupting natural breathing patterns. More details of olfactometer construction and usage are provided in the Supplemental Materials, along with detailed documentation in our open-source Github repository.

## 4 Data Processing and Analysis

Standard EEG preprocessing steps were applied: spherical spline interpolation of bad channels, average rereferencing, 0.5 Hz–100 Hz bandpass filtering and a 60 Hz notch filter, and removal of ocular artifacts via electrooculogram (EOG) regression using virtual EOG channels. The EEG data were recorded at 500 Hz and downsampled to 250 Hz. Epochs were baseline-corrected using a 1 s pre-stimulus window.

Nasal respiration data were recorded at the BME280 sensor’s native sampling rate of 8.76 Hz. Respiration signals were bandpass filtered at 0.1 Hz–0.4 Hz to isolate the respiratory rhythm and then upsampled to match the EEG sampling rate (250 Hz). Respiration was epoched using the same trial boundaries as EEG (TTL-locked; −1 s to 15 s relative to stimulus onset). Objective quality-control criteria were applied to exclude respiration trials affected by movement artifacts, sensor noise, or physiologically impossible breathing patterns. Specifically, trials were automatically removed if: (1) the respiration trace was near-flat (SD < 10^*−*3^), indicating sensor disconnection or artifact; (2) fewer than three respiratory cycles were detected within the 16 s epoch (peak prominence 0.05); or (3) no inhalation peak could be identified within − 0.4 s to 3 s relative to stimulus onset. These criteria ensured that only trials with reliable odor delivery timing and clean respiratory signals were analyzed. Overall, ∼80 % of trials were retained.

Frontal and temporal EEG channels were analyzed as our regions-of-interest, due to extensive literature demonstrating their involvement in human olfaction and respiration. Specifically, frontal channels target the orbitofrontal cortex, crucial for conscious odor perception, identification, and hedonic evaluation [35]. Temporal channels may capture projections from the primary olfactory cortex (piriform cortex), amygdala, and hippocampus, which are fundamental for initial odor processing, emotional responses, and memory formation [36]. These regions also exhibit respiratory rhythm entrainment, influencing olfactory processing [11, 37]. All reported p-values were corrected for multiple comparisons using Benjamini-Hochberg false discovery rate (FDR) correction unless otherwise specified.

### 4.1 Phase Amplitude Coupling

Recent studies have illuminated the connection between respiratory rhythms and neural oscillations, with accumulating evidence that nasal respiration modulates brain activity through direct mechanosensory pathways originating in the olfactory epithelium [11, 12, 38, 39]. Prior work has demonstrated respiratory modulation of cortical rhythms across multiple frequency bands [11, 40, 41], with implications for memory, attention, and sensory processing [12].

Given that our experimental paradigm involved olfactory stimulation during natural nasal breathing, we first sought to replicate and extend these foundational findings by characterizing respiratory-EEG coupling patterns in our participants. This analysis served two key purposes: (1) to verify respiration-locked modulation in the scalp EEG during natural breathing, and (2) to describe its frequency- and scalp-topographic distribution to contextualize subsequent odor-evoked analyses.

To compute respiratory-EEG coupling, continuous EEG and respiratory recordings (∼30 min per participant) were analyzed, which yielded approximately 300-450 breaths per subject. EEG data were bandpass filtered into frequency bands of interest: theta (4-8 Hz), alpha (8-12 Hz), beta (12-30 Hz), and low gamma (30-40 Hz). Amplitude envelopes were then extracted using the Hilbert transform. For the respiratory signal, phase was not derived via the Hilbert transform because respiration is not sinusoidal and inhale/exhale durations vary. Instead, respiratory phase was defined using cycle landmarks (peak/trough) and linearly interpolated such that 0^*°*^ corresponded to inhalation onset, 180^*°*^ to exhalation onset, and 360^*°*^ to the subsequent inhalation onset.

To quantify this respiratory-EEG synchrony, we employed phase-amplitude coupling (PAC), which measures how the phase of a lower-frequency signal (respiration) modulates the amplitude of a higher-frequency signal (EEG oscillations). Specifically, we used the modulation index (MI) [42], which quantifies the degree to which EEG amplitudes cluster at specific respiratory phases rather than distributing uniformly across the respiratory cycle. Higher MI values indicate stronger phase-amplitude coupling, with MI = 0 indicating no coupling (uniform distribution). For each breath, MI was computed per channel and frequency band, using respiratory phase as the modulating signal and the band-limited EEG amplitude envelope as the modulated signal.

#### Statistical Testing

Statistical significance was assessed using a surrogate data approach with 1000 iterations. Surrogate datasets were generated by circularly shifting the EEG amplitude time series relative to the respiratory phase by a random amount (minimum 30 seconds, equivalent to ∼3 respiratory cycles at the slowest breathing rate). This temporal shuffling approach disrupted respiratory–EEG coupling while maintaining the autocorrelation structure and statistical properties of both signals [43, 44]. Whole-head coupling strength was first summarized within each subject by averaging MI across channels and converting to a subject-level *z*-score relative to that subject’s surrogate distribution.

#### Spatial cluster analysis of PAC

To characterize the scalp topography of respiration–EEG PAC, we tested whether empirical MI significantly differed from chance-level MI at the group level using a cluster-based permutation procedure [44, 45]. This method identified significant differences across sensors while controlling for multiple comparisons. For each frequency band, empirical MI was compared to chance-level MI across participants using a one-sample t-test at each sensor. Candidate clusters were defined as spatially contiguous sensors (based on an EEG adjacency graph) exceeding a t-threshold of *p* < 0.05 (one-sided). Each cluster was characterized by the sum of t-values across its constituent sensors. Cluster significance was assessed by comparing observed cluster statistics to a null distribution generated from 1000 permutations, where participant z-scores were randomly sign-flipped.

### 4.2 Respiratory Phase-Locked Event-Related Spectral Perturbations (RPL-ERSPs)

Event-related potentials (ERPs) have long been considered the gold standard for analyzing neural responses to discrete events. However, ERPs are inherently limited by averaging time-domain waveforms across trials, which primarily preserves evoked activity — components with consistent timing relative to stimulus onset [46]. This approach necessarily discards induced activity, which is often more critical for understanding complex sensory and cognitive processing [47].

Event-related spectral perturbations (ERSPs) overcome this limitation by quantifying time- and frequency-specific changes in spectral power, thereby capturing both evoked and induced oscillatory dynamics [48, 49]. Induced activity, frequently observed in the theta, beta, and gamma frequency ranges, reflects ongoing brain processes such as sensory integration, attention, and cross-modal coordination, which may not align precisely in phase with the stimulus [50, 51].

This distinction is particularly crucial for brain–body coupling paradigms involving respiration, where breathing rhythms modulate cortical oscillatory activity in a frequency- and region-specific manner [38]. Respiration-locked oscillations in the piriform cortex and hippocampus contain both evoked components tightly linked to inhalation onset and induced power changes reflecting extended, odor-specific processing across sniff cycles [11, 52]. By preserving both response types, ERSPs enable detection of behaviorally relevant oscillatory modulations that ERPs alone may overlook.

Despite these advantages, traditional ERSPs calculated as frequency-over-time responses present challenges in respiration-gated paradigms. While aligning trials to inhalation onset at t=0 is common, respiratory cycle durations can vary substantially both within and across subjects (e.g., from approximately 2–6 s). As a result, group-averaged time-domain ERSPs implicitly assume consistent temporal alignment of inhale/exhale dynamics and their associated neural responses across trials, an assumption that may obscure respiration-locked spectral structure. To address this limitation, we introduce Respiratory Phase-Locked ERSPs (RPL-ERSPs), which represent oscillatory activity as a function of normalized respiratory phase rather than absolute time. Prior studies have demonstrated respiration-locked neural dynamics by aligning activity to respiratory events such as inhalation [11, 53]; however, because these time-locked analyses preserve the absolute duration of each breath, averaging across trials conflates different respiratory phases whenever cycle durations differ. RPL-ERSPs address this by mapping each respiratory cycle onto a normalized phase representation consisting of 360 discrete phase bins 1°-width, spanning the full breath cycle (0° = inhalation onset, 180° = exhalation onset, 360° = subsequent inhalation onset), enabling phase-resolved spectral analysis independent of absolute respiratory duration (Fig. 3**A**). Each phase bin aggregates all EEG samples whose respiratory phase falls within that bin, and spectral power is computed using the continuous Morlet wavelet transform. For our analysis, RPL-ERSPs were calculated over the first full breath after stimulus presentation for all trials, conditions, and subjects. This framework permits direct comparison of oscillatory dynamics across breaths and subjects despite substantial variability in breathing rate.

**Figure 3.**
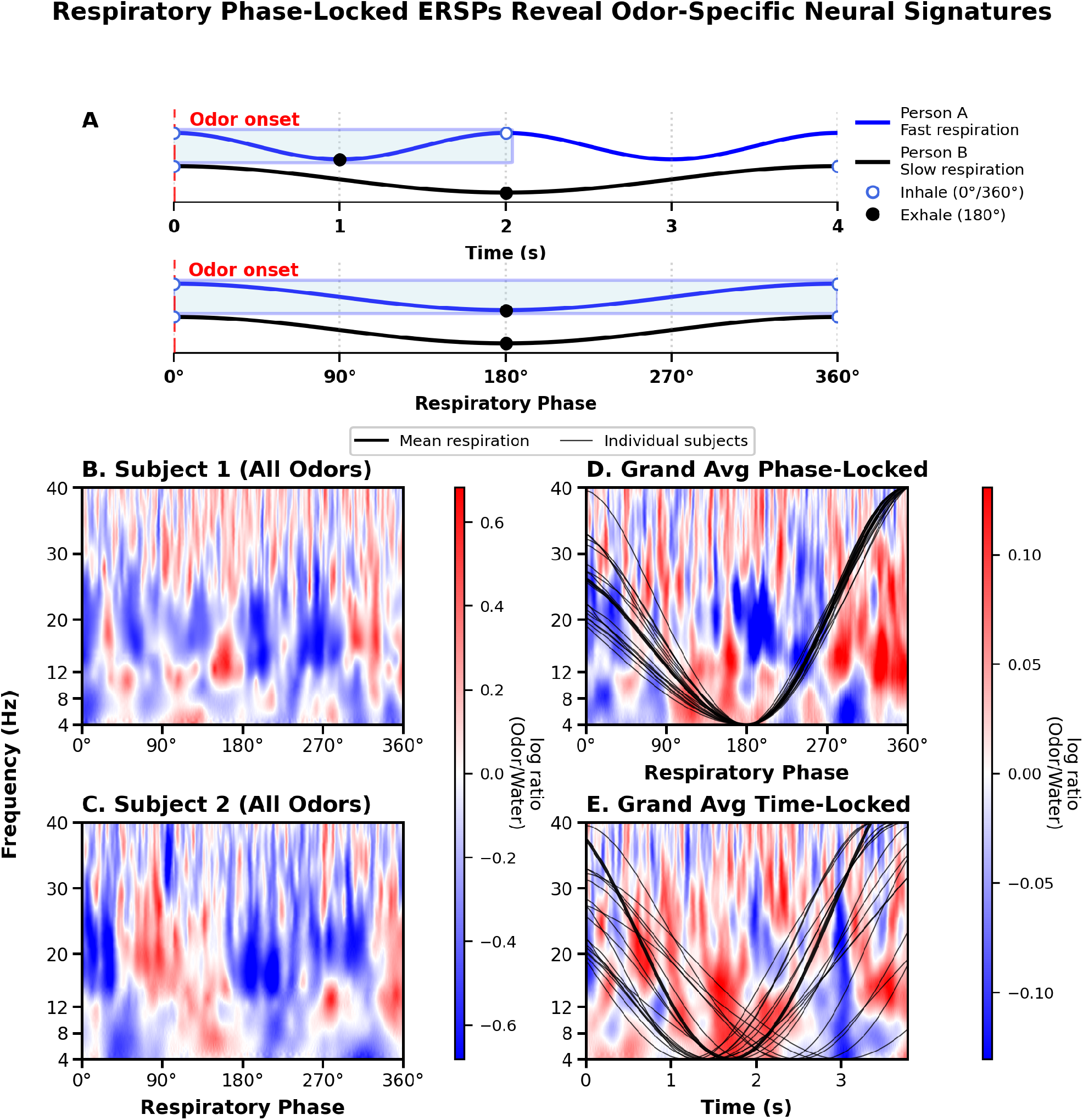
Respiratory phase-locked event-related spectral perturbations (RPL-ERSPs) characterize olfactory-evoked dynamics across the respiratory cycle. **A:** Conceptual schematic illustrating phase-locked analysis, which aligns neural activity to respiratory phase (0° = inhale onset; 180° = exhale onset) rather than absolute time. **B–C:** Representative single-subject RPL-ERSPs (averaged across odors) demonstrating individual-level phase-locked spectral structure across the full breath cycle. **D:** Grand-average RPL-ERSP across subjects (*n* = 17; averaged across odors) with overlaid respiratory waveforms (thin black traces: individual subjects; thick black: group mean). Phase alignment reduces between-subject variability in breath duration and sharpens phase-specific spectral features. **E:** Grand-average traditional time-locked ERSP computed from the same data window, shown with the same overlaid respiratory waveforms. Inter-subject variability in breath timing produces temporal smearing and reduced spectral contrast relative to the phase-locked representation.

The continuous Morlet wavelet is defined as:

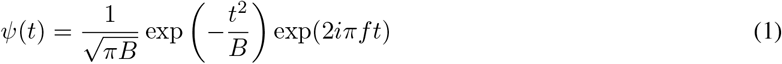

where *B* is the bandwidth, determining the temporal spread of the wavelet, and *f* is the center frequency, corresponding to the specific oscillatory frequency being analyzed. The Morlet wavelet closely mimics the expected oscillatory pattern in EEG signals, making it an ideal choice for time-frequency analysis [54]. A complex Morlet wavelet (PyWavelets [55], with bandwidth parameter *B* = 1.0) was used, which provides a practical balance between temporal and spectral resolution.

### 4.3 Classification Analysis

#### Feature Extraction

To assess whether RPL-ERSP features could discriminate odor from odorless control presentations, we compared our phase-locked approach to traditional time-locked approaches using early trial data (trials 1-3) from 17 normosmic participants. Early trials were selected to minimize habituation effects while providing sufficient data for stable spectral estimates. For each participant and condition, ERSPs were averaged across the first three trials to create subject-level representations.

Features comprised mean log-transformed spectral power within three frequency bands (theta: 4-8 Hz, alpha: 8-12 Hz, beta: 12-30 Hz). For RPL-ERSPs, the respiratory cycle was divided into four equal phase bins (0-90°, 90-180°, 180-270°, 270-360°) capturing early inhale, late inhale, early exhale, and late exhale, yielding 12 candidate features per sample (3 bands × 4 phase bins). For traditional time-locked ERSPs, we systematically optimized time window binning by testing three strategies over a window matched to average breath duration (3.78s): 1-bin (full window), 2-bin (half-duration bins), and 4-bin (quarter-duration bins). For each odor, we selected the binning strategy yielding the highest cross-validated AUC, ensuring time-locked ERSPs were evaluated under optimal conditions.

#### Nested Cross-Validation and Model Training

We implemented nested leave-one-subject-out (LOSO) cross-validation with two loops: an outer loop for performance evaluation and an inner loop for feature selection. In each outer fold, one subject’s data were held out while the remaining subjects formed the training set. We evaluated all possible feature subset sizes (*k* = 1 to 12), selecting the optimal *k* that maximized inner-loop area under the ROC curve (AUC). A final *𝓁*_2_ regularized logistic regression classifier was trained on the outer training set using only the selected features and evaluated on the held-out subject. Features were z-scored to prevent information leakage.

Performance was quantified using AUC from aggregated predictions across all outer folds. Statistical significance was assessed via permutation testing (1000 iterations), randomly shuffling labels within subjects and re-executing the nested LOSO procedure to generate null distributions. To identify features that were consistently informative, we ranked features by the average absolute value of their logistic-regression weights across the inner LOSO folds, so that features with consistently large weights were considered more stable and more predictive.

### 4.4 Free Response Odor Identification

To quantitatively score naturalistic odor descriptions, a hybrid computational approach was implemented that combined empirical odor databases with modern natural language processing. Professional odor descriptors were extracted from the GoodScents database [56] and, for each target odor, a set of chemically related odorants (e.g., menthol, menthone, peppermint oil) was assembled. Descriptor frequencies were then tallied across these odor-relevant GoodScents entries and converted into association weights (the fraction of entries containing each descriptor), yielding a descriptor profile that reflected how strongly each term was linked to the target odor in expert annotations.

For semantic matching, we employed a sentence-transformer model (all-mpnet-base-v2) [18] that mapped text into high-dimensional vector spaces where semantically related terms cluster together. We computed cosine similarity between these word embeddings to generate continuous similarity scores rather than binary matches.

Final identification scores combined both information sources equally:

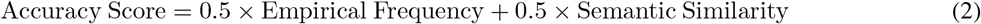

### 4.5 Olfactory Habituation

Olfactory habituation – the progressive reduction in neural and perceptual response to repeated odor exposures – is a fundamental property of chemosensory systems that reflects the brain’s ability to filter redundant stimuli and prioritize novel inputs [57]. EEG studies have documented amplitude reductions in ERPs with repeated exposure [58, 59], though the functional consequences for odor discriminability remain poorly understood. Altered habituation patterns have been documented in Parkinson’s disease [60] and autism spectrum disorders [61], suggesting that quantifying adaptation dynamics may provide clinically valuable biomarkers. Understanding how olfactory habituation affects phase-locked neural signatures is therefore both theoretically important for models of sensory adaptation and practically relevant for developing passive diagnostic tools.

To directly quantify changes in neural discriminability, we applied the nested LOSO classification framework described above, separately to early trials (1-3) and late trials (6-8). We hypothesized that habituation would manifest as: (1) decreased AUC from early to late trials, and (2) shifts in feature selection patterns, particularly affecting the most discriminative phase-locked features identified in early trials.

## 5 Results

### 5.1 Phase Amplitude Coupling

#### Overall PAC Strength

Whole-head respiration–EEG PAC was significant in the alpha, beta, and low-gamma bands (*n* = 17 subjects; Fig. 2A): alpha (*p* = 0.043, Cohen’s *d* = 0.48), beta (*p* = 0.043, *d* = 0.50), and low gamma (*p* = 0.011, *d* = 0.80). Theta-band coupling was not significant (*p* = 0.232, *d* = 0.19).

#### Spatial Specificity

Cluster-based permutation testing revealed band-specific scalp distributions (Fig. 2B). Row B shows 2D topographic maps of the channel-wise test statistics, with significant cluster electrodes marked. In the alpha band, two clusters were observed: a fronto-temporal cluster (*p* = 0.022, *n* = 23 channels) and a parietal-temporal cluster (*p* = 0.040, *n* = 18 channels). In the low-gamma band, PAC formed a broad fronto-temporal cluster (*p* = 0.001, *n* = 48 channels), consistent with prior reports of respiration-gamma coupling over frontal, frontocentral, and temporoparietal regions [62]. No significant spatial clusters were detected for theta or beta bands.

### 5.2 Respiratory Phase-Locked ERSPs

Inspection of the grand-averaged RPL-ERSP (Fig. 3**D**, pooled across odors) revealed two consistent features. Theta-band power increased during late inhalation, consistent with prior reports of increased theta activity during active olfactory sampling [11, 13]. Alpha-beta power (8–20 Hz) also decreased during exhalation—a pattern not previously reported, as human olfactory research has focused almost exclusively on inhalation when odor enters the nasal cavity. This exhalation-locked suppression may indicate continued odor processing during the non-sampling phase: alpha decreases are associated with attentional engagement [63], while beta modulation has been linked to learning-dependent reorganization of olfactory representations [64].

Comparison of phase-locked (Fig.3 **D**) versus time-locked (Fig.3**E**) ERSPs computed from identical data demonstrates the advantage of phase alignment in respiration-gated paradigms. Overlaid black curves show individual subjects’ respiratory traces, revealing substantial inter-subject variability in breath duration (∼2 s-6 s). In the time-locked ERSP (Fig.3**E**), this variability causes neural activity from different respiratory phases to average together, producing temporal smearing and reduced spectral contrast. Phase-locked alignment (Fig.3**D**) yields sharper delineation of theta enhancement during inhalation and alpha-beta suppression during exhalation, demonstrating that respiratory phase provides a more appropriate reference frame than absolute time when breath timing varies across individuals.

Although there is more variability, representative single-subject examples are shown in Fig. 3**B–C**, illustrating that these phase-locked spectral features are observable at the individual-subject level. Group averaged RPL-ERSPs for specific odors are also shown in Supplemental Fig.S1.

### 5.3 Classification Performance

Despite systematic optimization of temporal binning strategies for each odor, RPL-ERSPs consistently outperformed time-locked approaches across all three odorants (Fig.4**B**) For peppermint vs. water, RPL-ERSP achieved AUC = 0.785, exceeding the best time-locked strategy (4-bin, AUC = 0.495) by ΔAUC = +0.290. Oregano vs. water classification showed a similar pattern, with RPL-ERSP (AUC = 0.671) outperforming the optimal time-locked configuration (4-bin, AUC = 0.595) by ΔAUC = +0.076. Grapefruit vs. water discrimination yielded the smallest performance gain, with RPL-ERSP (AUC = 0.617) surpassing optimized time-locked ERSP (4-bin, AUC = 0.613) by ΔAUC = +0.005 (Fig.4**B**). Critically, these results demonstrate that the advantage of respiratory phase alignment cannot be attributed to suboptimal temporal binning in time-locked analyses, nor to differences in analysis window duration.

#### Feature Selection Patterns

Analysis of feature selection frequency across outer LOSO folds revealed distinct patterns between phase-locked and time-locked approaches (Fig. 4**A**). For RPL-ERSPs, alpha-band activity during exhalation phases showed the strongest and most consistent selection: features from 180-270° and 270-360° bins were selected in 100% of folds across all odors, indicating a robust, odor-general neural signature during the alpha band exhalation phase. Theta-band features were primarily selected during the inhalation period with 0-90° bins selected in 64.7% of folds, and 90-180° selected in 87.7% of folds on average. Beta-band activity during exhalation showed modest but focused phase-specificity during exhalation. Across outer LOSO folds, an average of 5.0 features were used.

**Figure 4.**
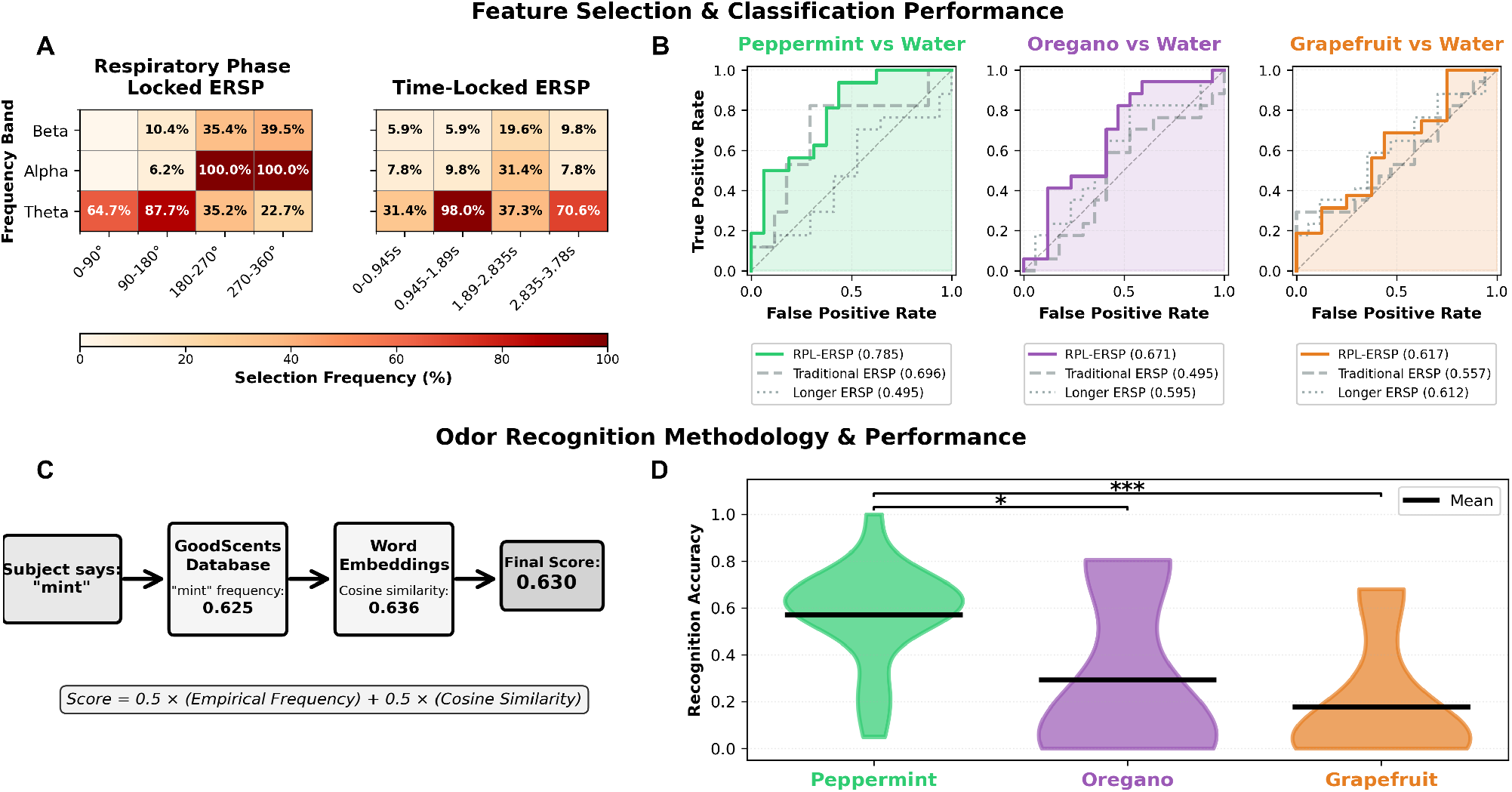
Respiratory phase-locked features improve odor classification and align with behavioral recognition performance. **A:** Feature selection frequencies across respiratory phase bins and frequency bands demonstrate that respiratory phase-locked ERSP (RPL-ERSP) features are preferentially selected by the classifier compared to traditional time-locked features. RPL-ERSPs show high selection rates for alpha-band activity during early exhalation (180°-270°, 64.7%) and late exhalation (270°-360°,100%), as well as theta-band activity during inhalation phases (0°-90°, 64.7%; 90°-180°, 87.7%). In contrast, traditional time-locked ERSP shows more distributed and lower selection frequencies, indicating less discriminative power. **B:** ROC curves confirm superior classification performance for RPL-ERSP across all odors. Peppermint vs water achieves highest discrimination (AUC=0.785), followed by oregano (AUC=0.671) and grapefruit (AUC=0.617). Traditional time-locked ERSPs with optimized binning show degraded performance (peppermint: 0.696, oregano: 0.495, grapefruit: 0.557), while extending the time window to match average breath duration (3.78s) provides minimal improvement. **C:** Odor recognition scoring methodology combined empirical descriptor frequencies from the GoodScents database with semantic word embeddings. For each subject response, we compute a hybrid score as the average of empirical frequency (how often the descriptor appears in perfumery databases for that odor) and cosine similarity between word embeddings. Example: “mint” for peppermint yields empirical frequency of 0.625 and semantic similarity of 0.636, resulting in final score of 0.630. **D:** Behavioral recognition accuracy parallels the neural classification odor hierarchy. Peppermint shows highest recognition (mean=0.57±0.26), significantly better than both oregano (mean=0.30±0.35, p<0.05) and grapefruit (mean=0.18±0.26, p<0.001). Oregano shows higher recognition than grapefruit. Individual subject variability shown in violin plots with horizontal lines indicating mean performance. Together, these results establish: (1) phase-locked neural features provide superior classification over traditional time-locked features and (2) behavioral recognition accuracy parallels neural discriminability.

In contrast, time-locked approaches showed fundamentally different patterns. Rather than identifying specific discriminative windows, time-locked ERSP showed broad theta dominance across all temporal bins, with minimal alpha and beta contributions, using an average of 3.4 features. This suggested that temporal averaging smeared odor-related dynamics across the entire breath, collapsing multi-band spectral structure into a single dominant frequency component. The ubiquitous but variable theta selection indicated that time-locking failed to capture the precise phase-dependent neural dynamics that characterized odor processing.

### 5.4 Free Response Odor Identification

Free-form identification accuracy varied significantly across odors, mirroring the hierarchy observed in EEG classification (Fig. 4**D**). Peppermint showed the highest identification accuracy (mean: 0.572 ± 0.213), significantly exceeding both oregano (mean: 0.295 ± 0.353) and grapefruit (mean: 0.178 ± 0.257).

This behavioral hierarchy (peppermint > oregano > grapefruit) directly parallels our EEG classification results in Fig. 4**B**. The convergence between implicit neural discriminability and explicit verbal identification provides crucial validation: stimuli eliciting more separable neural representations also yield more accurate naming, suggesting that EEG patterns reflect genuine perceptual differences rather than low-level confounds.

### 5.5 Olfactory Habituation

Early-trial classification (trials 1-3) achieved strong odor-water discrimination (mean AUC = 0.691 ± 0.070; peppermint: 0.785, oregano: 0.671, grapefruit: 0.617). By late trials (6-8), performance declined to near-chance levels (mean AUC = 0.427 ± 0.024)—a 38% reduction that was highly significant (paired t-test: t(2) = 8.11, p = 0.007), indicating neural responses became indistinguishable from water baseline (Fig. 5**A**).

**Figure 5.**
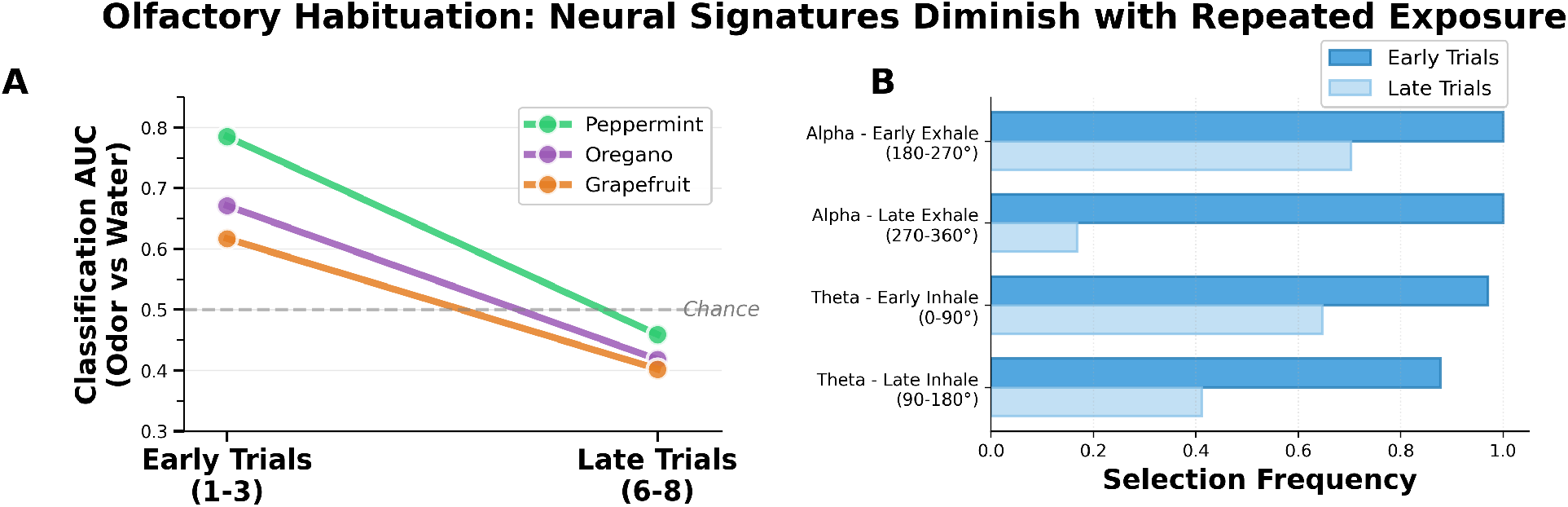
Olfactory neural signatures decline with repeated exposure, demonstrating rapid adaptation across odors and respiratory phases. **A:** Classification performance (AUC) for discriminating each odor from water declines significantly from early to late trial blocks across all odors. Peppermint maintains highest discrimination throughout (early: 0.785, late: 0.461, Δ=-41%), followed by oregano (early: 0.671, late: 0.454, Δ=-32%) and grapefruit (early: 0.617, late: 0.403, Δ=-35%). All odors approach chance performance by late trials, indicating substantial neural adaptation. **B:** Feature selection frequencies reveal which respiratory phase bins drive classification and how they change with habituation. Early trials show robust reliance on alpha band during exhale, as well as theta band during inhale. Late trials show dramatic declines in feature importance: alpha early exhale drops by 30%, alpha late exhale by 83%, theta early inhale by 33%, and theta late inhale by 53%. Compensatory recruitment of beta-band features fails to restore discriminability. Together, these results establish rapid neural habituation to repeated odor presentations, with preferential adaptation of the neural activity that initially provided the strongest discriminative information.

This classification decline was accompanied by systematic changes in feature selection patterns (Fig.5**B**). In early trials, the classifier robustly selected alpha-band exhalation and theta-band inhalation power across 88-100% of LOSO folds. In late trials, these discriminative features declined dramatically: alpha activity during late exhalation (270-360°) dropped by 83%, while theta activity during late inhalation (90-180°) dropped by 53%. The classifier attempted to compensate by recruiting beta-band activity during early inhalation (increasing from 0% to 83% selection frequency), yet this failed to restore discriminability, with AUC remaining near chance.

## 6 Discussion

Through multiple lines of evidence, this study demonstrates that natural respiration is a meaningful reference frame for odor-evoked cortical dynamics. First, building on prior work showing respiration–EEG phase-amplitude coupling in alpha and gamma bands [62], we confirmed significant coupling in our dataset (alpha, beta, low-gamma), though theta PAC was absent—consistent with scalp EEG’s limited sensitivity to deep medial temporal sources where theta coupling has been observed intracranially [11]. To capture respiration-EEG interactions, we then introduced respiratory phase–locked event-related spectral perturbations (RPL-ERSPs). This phase-based framework avoids a limitation of conventional time-locking, where natural variability in breath duration can smear or hide respiratory phase-locked responses. Accordingly, phase-locked features outperformed optimized time-locked analyses across windows and frequency bands, driven by multi-band, respiratory phase-specific signatures—especially theta increases (4–8 Hz) during inhalation and alpha decreases (8–12 Hz) during exhalation. The theta-inhalation pattern is consistent with intracranial findings showing theta oscillations during inhalation enhance odor processing [13], demonstrating that these dynamics can be captured at the scalp level using RPL-ERSPs. The alpha-exhalation signature has not been previously documented, as human olfactory research has focused almost exclusively on inhalation when odor enters the nasal cavity. Together, these complementary patterns may serve as neural markers of odor processing across the full respiratory cycle. The consistency of these signatures in healthy participants suggests that they could form the basis for objective clinical assessment: quantifying individual deviations from this respiratory phase-locked pattern may offer a non-invasive, scalp-based approach to detecting and characterizing olfactory dysfunction.

RPL-ERSPs also provided a sensitive window into habituation: neural discriminability dropped markedly across repetitions, with mean AUC declining from 0.69 in early trials to near-chance (0.43) after just ∼6–8 exposures. The most informative features (alpha-exhalation and theta-inhalation) showed the largest reductions in selection frequency (up to 83% decline). This may suggest a reorganization of neural representations that’s more complex than amplitude attenuation alone, extending beyond the chemosensory event-related potential reductions documented in prior studies [58, 59] which primarily capture evoked activity. The ERSP approach used here preserves both evoked and induced oscillatory activity, raising the possibility that olfactory habituation and neural discriminability depend not only on evoked responses but also on reorganization of induced oscillatory dynamics. Measuring this decline in neural discriminability may prove clinically valuable, as olfactory habituation rates differ systematically in neurological populations, including Parkinson’s disease [60] and autism [61]. The rate at which classification performance drops from above-chance to baseline may itself vary across populations, offering a quantitative marker of atypical habituation. By quantifying adaptation dynamics passively without task engagement, these oscillatory features offer a potential tool for early detection and phenotyping in populations where habituation mechanisms are disrupted.

In parallel, our free-response scoring methodology combined empirical odor descriptor databases with word embeddings from sentence-transformer models, offering a promising alternative to forced-choice paradigms. Because odor knowledge is often communicated in flexible, context-dependent language rather than fixed labels, participants may express recognition through associations (“what it reminds me of”) or broader categorical descriptors (e.g., peppermint as “gum,” oregano as “herbal”) even when the canonical name is unavailable. By scoring responses in semantic space, our approach captures this graded, naturalistic form of recognition instead of treating it as simply incorrect, potentially improving sensitivity in clinical populations where explicit naming is impaired but perceptual recognition remains partially intact. Notably, the neural discriminability of odors (peppermint > oregano > grapefruit) mirrored participants’ behavioral identification accuracy from free-form descriptions, demonstrating that RPL-ERSPs capture perceptually meaningful distinctions rather than random neural variance. This framework opens promising avenues for integrating language-based odor representations with computational models of olfactory perception and neural activity, potentially enabling personalized odor embeddings that capture individual differences in emotion and memory associations beyond population-averaged responses.

These findings were enabled by designing an open-source, respiration-synchronized olfactometer that used real-time respiratory monitoring to automate odor delivery at a fraction of the cost of comparable systems (<$500 vs. >$30,000 [14]). This platform provides an affordable way to measure olfactory processing during natural respiration, avoiding the forced inhalation paradigms used in prior work. We anticipate that this respiration-locked, low-cost platform will be especially valuable in settings where conventional olfactory tests are impractical or systematically biased by cognitive, linguistic, or motor demands (e.g., Parkinson’s, Alzheimer’s, Autism Spectrum Disorder), enabling objective assessment without sustained task engagement.

Our findings establish that natural respiratory phase provides a robust organizing principle for olfactory cortical dynamics. We identify consistent, phase-specific signatures across the full respiratory cycle—theta increases during inhalation and alpha decreases during exhalation—captured non-invasively through affordable instrumentation. By integrating language-flexible behavioral assessment with these phase-locked patterns, this framework offers a scalable platform for both basic olfactory neuroscience and clinical translation—enabling objective assessment in populations where conventional testing is impractical or biased. Future work should validate these findings in larger and more diverse samples spanning age ranges, clinical populations with olfactory dysfunction, and cultural contexts, and expand beyond three odors to assess generalization across the olfactory stimulus space.

## 6.0.1 Acknowledgments

This work was supported by the National Science Foundation Graduate Research Fellowship Program (DGE-2146755), Wu Tsai Neurosciences Institute, Koret Human Neurosciences Community Laboratory, and Stanford University Bio-X Fellowship. We would also like to thank Syamantak Payra for his help with circuitry.

The authors declare that they have no competing interests.

## References

[1] Xiuli Dan et al. “Olfactory dysfunction in aging and neurodegenerative diseases”. In: Ageing Research Reviews 70 (Sept. 2021), p. 101416. ISSN: 1568-1637. DOI: 10.1016/j.arr.2021.101416. URL: http://dx.doi.org/10.1016/j.arr.2021.101416.

[2] Christopher Hawkes. “Olfaction in Neurodegenerative Disorder”. In: Taste and Smell. KARGER, 2006, pp. 133–151. ISBN: 3805581238. DOI: 10.1159/000093759. URL: http://dx.doi.org/10.1159/000093759.

[3] Concepció Marin et al. “Olfactory Dysfunction in Neurodegenerative Diseases”. In: Current Allergy and Asthma Reports 18.8 (June 2018). ISSN: 1534-6315. DOI:10.1007/s11882-018-0796-4. URL: http://dx.doi.org/10.1007/s11882-018-0796-4.

[4] Woodward et al. “Validation of olfactory deficit as a biomarker of Alzheimer disease”. In: Neurology Clinical Practice 7.1 (Feb. 2017), pp. 5–14. ISSN: 2163-0933. DOI:10.1212/cpj.0000000000000293. URL: http://dx.doi.org/10.1212/CPJ.0000000000000293.

[5] Richard L. Doty et al. “University of pennsylvania smell identification test: A rapid quantitative olfactory function test for the clinic”. In: The Laryngoscope 94.2 (Feb. 1984), pp. 176–178. ISSN: 1531-4995. DOI: 10.1288/00005537-198402000-00004. URL: http://dx.doi.org/10.1288/00005537-198402000-00004.

[6] B. N. Johnson et al. “A Comparison of Methods for Sniff Measurement Concurrent with Olfactory Tasks in Humans”. In: Chemical Senses 31.9 (Aug. 2006), pp. 795–806. ISSN: 1464-3553. DOI:10.1093/chemse/bjl021. URL: http://dx.doi.org/10.1093/chemse/bjl021.

[7] Marcel S. Kehl et al. “Single-neuron representations of odours in the human brain”. In: Nature 634.8034 (Oct. 2024), pp. 626–634. ISSN: 1476-4687. DOI: 10.1038/s41586-024-08016-5. URL:http://dx.doi.org/10.1038/s41586-024-08016-5.

[8] Artin Arshamian et al. “Respiration Modulates Olfactory Memory Consolidation in Humans”. In: The Journal of Neuroscience 38.48 (Oct. 2018), pp. 10286–10294. ISSN: 1529-2401. DOI: 10.1523/jneurosci.3360-17.2018. URL: http://dx.doi.org/10.1523/JNEUROSCI.3360-17.2018.

[9] Roman Shusterman et al. “Precise olfactory responses tile the sniff cycle”. In: Nature Neuroscience 14.8 (July 2011), pp. 1039–1044. ISSN: 1546-1726. DOI: 10.1038/nn.2877. URL: http://dx.doi.org/10.1038/nn.2877.

[10] Keiji Miura, Zachary F. Mainen, and Naoshige Uchida. “Odor Representations in Olfactory Cortex: Distributed Rate Coding and Decorrelated Population Activity”. In: Neuron 74.6 (June 2012), pp. 1087–1098. ISSN: 0896-6273. DOI: 10.1016/j.neuron.2012.04.021. URL: http://dx.doi.org/10.1016/j.neuron.2012.04.021.

[11] Christina Zelano et al. “Nasal Respiration Entrains Human Limbic Oscillations and Modulates Cognitive Function”. In: The Journal of Neuroscience 36.49 (Dec. 2016), pp. 12448–12467. ISSN: 1529-2401. DOI: 10.1523/jneurosci.2586-16.2016. URL: http://dx.doi.org/10.1523/JNEUROSCI.2586-16.2016.

[12] Adriano B.L. Tort, Jurij Brankacč k, and Andreas Draguhn. “Respiration-Entrained Brain Rhythms Are Global but Often Overlooked”. In: Trends in Neurosciences 41.4 (Apr. 2018), pp. 186–197. ISSN: 0166-2236. DOI: 10.1016/j.tins.2018.01.007. URL: http://dx.doi.org/10.1016/j.tins.2018.01.007.

[13] Heidi Jiang et al. “Theta Oscillations Rapidly Convey Odor-Specific Content in Human Piriform Cortex”. In: Neuron 94.1 (Apr. 2017), 207–219.e4. ISSN: 0896-6273. DOI: 10.1016/j.neuron.2017.03.021. URL: http://dx.doi.org/10.1016/j.neuron.2017.03.021.

[14] Tyler S. Lorig et al. “A computer-controlled olfactometer for fMRI and electrophysiological studies of olfaction”. In: Behavior Research Methods, Instruments, & Computers 31.2 (June 1999), pp. 370–375. ISSN: 1532-5970. DOI: 10.3758/bf03207734. URL: http://dx.doi.org/10.3758/BF03207734.

[15] Margareta Hedner et al. “Cognitive factors in odor detection, odor discrimination, and odor identification tasks”. In: Journal of Clinical and Experimental Neuropsychology 32.10 (Apr. 2010), pp. 1062–1067. ISSN: 1744-411X. DOI: 10.1080/13803391003683070. URL: http://dx.doi.org/10.1080/13803391003683070.

[16] Jonas K. Olofsson and Jay A. Gottfried. “The muted sense: neurocognitive limitations of olfactory language”. In: Trends in Cognitive Sciences 19.6 (June 2015), pp. 314–321. ISSN: 1364-6613. DOI: 10.1016/j.tics.2015.04.007. URL: http://dx.doi.org/10.1016/j.tics.2015.04.007.

[17] William S. Cain. “To Know with the Nose: Keys to Odor Identification”. In: Science 203.4379 (Feb. 1979), pp. 467–470. ISSN: 1095-9203. DOI: 10.1126/science.760202. URL: http://dx.doi.org/10.1126/science.760202.

[18] Nils Reimers and Iryna Gurevych. “Sentence-BERT: Sentence Embeddings using Siamese BERT-Networks”. In: Proceedings of the 2019 Conference on Empirical Methods in Natural Language Processing. Association for Computational Linguistics, Nov. 2019. URL: https://arxiv.org/abs/1908.10084.

[19] Richard L. Doty and Vidyulata Kamath. “The influences of age on olfaction: a review”. In: Frontiers in Psychology 5 (2014). ISSN: 1664-1078. DOI: 10.3389/fpsyg.2014.00020. URL: http://dx.doi.org/10.3389/fpsyg.2014.00020.

[20] Asifa Majid. “Human Olfaction at the Intersection of Language, Culture, and Biology”. In: Trends in Cognitive Sciences 25.2 (Feb. 2021), pp. 111–123. ISSN: 1364-6613. DOI: 10.1016/j.tics.2020.11.005. URL: http://dx.doi.org/10.1016/j.tics.2020.11.005.

[21] J. N. Lundstrom et al. “Olfactory Event-Related Potentials Reflect Individual Differences in Odor Valence Perception”. In: Chemical Senses 31.8 (2006), pp. 705–711. ISSN: 1464-3553. DOI: 10.1093/chemse/bjl012. URL: http://dx.doi.org/10.1093/chemse/bjl012.

[22] Manasweeta Angane et al. “Essential Oils and Their Major Components: An Updated Review on Antimicrobial Activities, Mechanism of Action and Their Potential Application in the Food Industry”. In: Foods 11.3 (Feb. 2022), p. 464. ISSN: 2304-8158. DOI: 10.3390/foods11030464. URL: http://dx.doi.org/10.3390/foods11030464.

[23] Fei Han et al. “Chemical composition and antioxidant activities of essential oils from different parts of the oregano”. In: Journal of Zhejiang University-SCIENCE B 18.1 (Jan. 2017), pp. 79–84. ISSN: 1862-1783. DOI: 10.1631/jzus.b1600377. URL: http://dx.doi.org/10.1631/jzus.B1600377.

[24] Avat Arman Taherpour et al. “Chemical composition analysis of the essential oil of Mentha piperita L. from Kermanshah, Iran by hydrodistillation and HS/SPME methods”. In: Journal of Analytical Science and Technology 8.1 (May 2017). ISSN: 2093-3371. DOI: 10.1186/s40543-017-0122-0. URL: http://dx.doi.org/10.1186/s40543-017-0122-0.

[25] David D. McKemy, Werner M. Neuhausser, and David Julius. “Identification of a cold receptor reveals a general role for TRP channels in thermosensation”. In: Nature 416.6876 (Feb. 2002), pp. 52–58. ISSN: 1476-4687. DOI:10.1038/nature719. URL: http://dx.doi.org/10.1038/nature719.

[26] The Good Scents Company. d-Limonene. https://www.thegoodscentscompany.com/data/rw1020691.html. Accessed: 2026. 2026.

[27] The Good Scents Company. Origanum Oil. https://www.thegoodscentscompany.com/odor/origanum.html. Accessed: 2026. 2026.

[28] doTERRA International. Essential Oils: Peppermint, Oregano, and Grapefruit. Certified Pure Tested Grade (CPTG). 2024.

[29] Spriek. Intelligent Digital Display Nebulizer. Commercial air compressor nebulizer. Available from Amazon. 2024.

[30] J. Lötsch et al. “Factors affecting pain intensity in a pain model based upon tonic intranasal stimulation in humans”. In: Inflammation Research 47.11 (Nov. 1998), pp. 446–450. ISSN: 1420-908X. DOI: 10.1007/s000110050359. URL: http://dx.doi.org/10.1007/s000110050359.

[31] Johan N. Lundström et al. “Methods for building an inexpensive computer-controlled olfactometer for temporally-precise experiments”. In: International Journal of Psychophysiology 78.2 (Nov. 2010), pp. 179–189. ISSN: 0167-8760. DOI: 10.1016/j.ijpsycho.2010.07.007. URL: http://dx.doi.org/10.1016/j.ijpsycho.2010.07.007.

[32] Brad Johnson, Rehan M. Khan, and Noam Sobel. “Human Olfactory Psychophysics”. In: The Senses: A Comprehensive Reference. Elsevier, 2008, pp. 823–857. ISBN: 9780123708809. DOI: 10.1016/b978-012370880-9.00130-4. URL: http://dx.doi.org/10.1016/B978-012370880-9.00130-4.

[33] S. Al Aïn and J.A. Frasnelli. “Intranasal Trigeminal Chemoreception”. In: Conn’s Translational Neuroscience. Elsevier, 2017, pp. 379–397. ISBN: 9780128023815. DOI: 10.1016/b978-0-12-802381-5.00030-0. URL: http://dx.doi.org/10.1016/B978-0-12-802381-5.00030-0.

[34] RoboticsBackEnd. Arduino Fast Digital Write. 2021. URL: https://roboticsbackend.com/arduino-fast-digitalwrite/#:~:text=We%20have%20the%20answer:%20a,have%20a%20much%20better%20precision..

[35] Edmund T. Rolls. “The functions of the orbitofrontal cortex”. In: Brain and Cognition 55.1 (June 2004), pp. 11–29. ISSN: 0278-2626. DOI: 10.1016/s0278-2626(03)00277-x. URL: http://dx.doi.org/10.1016/S0278-2626(03)00277-X.

[36] A.M. Lascano et al. “Spatio–temporal dynamics of olfactory processing in the human brain: an event-related source imaging study”. In: Neuroscience 167.3 (May 2010), pp. 700–708. ISSN: 0306-4522. DOI: 10.1016/j.neuroscience.2010.02.013. URL: http://dx.doi.org/10.1016/j.neuroscience.2010.02.013.

[37] Brian Dlouhy et al. “Human forebrain neural synchronization and entrainment to breathing during wakefulness, sleep, and external mechanical ventilation”. In: (May 2025). DOI: 10.21203/rs.3.rs-6568046/v1. URL: http://dx.doi.org/10.21203/rs.3.rs-6568046/v1.

[38] Daniel S. Kluger and Joachim Gross. “Respiration modulates oscillatory neural network activity at rest”. In: PLOS Biology 19.11 (Nov. 2021). Ed. by David Poeppel, e3001457. ISSN: 1545-7885. DOI: 10.1371/journal.pbio.3001457. URL: http://dx.doi.org/10.1371/journal.pbio.3001457.

[39] Sadeq Mohammadi, Gholam-Ali Hossein-Zadeh, and Mohammad Reza Raoufy. “Breathing mode selectively modulates brain-wide functional connectivity”. In: PLOS One 20.11 (Nov. 2025). Ed. by Federico Giove, e0334165. ISSN: 1932-6203. DOI: 10.1371/journal.pone.0334165. URL: http://dx.doi.org/10.1371/journal.pone.0334165.

[40] Jose L. Herrero et al. “Breathing above the brain stem: volitional control and attentional modulation in humans”. In: Journal of Neurophysiology 119.1 (Jan. 2018), pp. 145–159. ISSN: 1522-1598. DOI: 10.1152/jn.00551.2017. URL: http://dx.doi.org/10.1152/jn.00551.2017.

[41] Ofer Perl et al. “Human non-olfactory cognition phase-locked with inhalation”. In: Nature Human Behaviour 3.5 (Mar. 2019), pp. 501–512. ISSN: 2397-3374. DOI: 10.1038/s41562-019-0556-z. URL: http://dx.doi.org/10.1038/s41562-019-0556-z.

[42] Adriano B. L. Tort et al. “Measuring Phase-Amplitude Coupling Between Neuronal Oscillations of Different Frequencies”. In: Journal of Neurophysiology 104.2 (Aug. 2010), pp. 1195–1210. ISSN: 1522-1598. DOI: 10.1152/jn.00106.2010. URL: http://dx.doi.org/10.1152/jn.00106.2010.

[43] Kurt E. Weaver et al. “Directional patterns of cross frequency phase and amplitude coupling within the resting state mimic patterns of fMRI functional connectivity”. In: NeuroImage 128 (Mar. 2016), pp. 238–251. ISSN: 1053-8119. DOI: 10.1016/j.neuroimage.2015.12.043. URL: http://dx.doi.org/10.1016/j.neuroimage.2015.12.043.

[44] Craig G. Richter et al. “Phase-amplitude coupling at the organism level: The amplitude of spontaneous alpha rhythm fluctuations varies with the phase of the infra-slow gastric basal rhythm”. In: NeuroImage 146 (Feb. 2017), pp. 951–958. ISSN: 1053-8119. DOI: 10.1016/j.neuroimage.2016.08.043. URL: http://dx.doi.org/10.1016/j.neuroimage.2016.08.043.

[45] Eric Maris and Robert Oostenveld. “Nonparametric statistical testing of EEG-and MEG-data”. In: Journal of Neuroscience Methods 164.1 (Aug. 2007), pp. 177–190. ISSN: 0165-0270. DOI:10.1016/j.jneumeth.2007.03.024. URL: http://dx.doi.org/10.1016/j.jneumeth.2007.03.024.

[46] Pasquale Arpaia et al. “Assessment and Scientific Progresses in the Analysis of Olfactory Evoked Potentials”. In: Bioengineering 9.6 (June 2022), p. 252. ISSN: 2306-5354. DOI: 10.3390/bioengineering9060252. URL: http://dx.doi.org/10.3390/bioengineering9060252.

[47] Caroline Huart et al. “Time-Frequency Analysis of Chemosensory Event-Related Potentials to Characterize the Cortical Representation of Odors in Humans”. In: PLoS ONE 7.3 (Mar. 2012). Ed. by Efthimios M. C. Skoulakis, e33221. ISSN: 1932-6203. DOI: 10.1371/journal.pone.0033221. URL: http://dx.doi.org/10.1371/journal.pone.0033221.

[48] Scott Makeig. “Auditory event-related dynamics of the EEG spectrum and effects of exposure to tones”. In: Electroencephalography and Clinical Neurophysiology 86.4 (Apr. 1993), pp. 283–293. ISSN: 0013-4694. DOI: 10.1016/0013-4694(93)90110-h. URL: http://dx.doi.org/10.1016/0013-4694(93)90110-H.

[49] Catherine Tallon-Baudry et al. “Oscillatory Gamma-Band (30–70 Hz) Activity Induced by a Visual Search Task in Humans”. In: The Journal of Neuroscience 17.2 (Jan. 1997), pp. 722–734. ISSN: 1529-2401. DOI: 10.1523/jneurosci.17-02-00722.1997. URL: http://dx.doi.org/10.1523/JNEUROSCI.17-02-00722.1997.

[50] Wolfgang Klimesch. “EEG alpha and theta oscillations reflect cognitive and memory performance: a review and analysis”. In: Brain Research Reviews 29.2–3 (Apr. 1999), pp. 169–195. ISSN: 0165-0173. DOI: 10.1016/s0165-0173(98)00056-3. URL: http://dx.doi.org/10.1016/s0165-0173(98)00056-3.

[51] Catherine Tallon-Baudry. “Oscillatory gamma activity in humans and its role in object representation”. In: Trends in Cognitive Sciences 3.4 (Apr. 1999), pp. 151–162. ISSN: 1364-6613. DOI: 10.1016/s1364-6613(99)01299-1. URL: http://dx.doi.org/10.1016/s1364-6613(99)01299-1.

[52] Nikolaos Karalis and Anton Sirota. “Breathing coordinates cortico-hippocampal dynamics in mice during offline states”. In: Nature Communications 13.1 (Jan. 2022). ISSN: 2041-1723. DOI: 10.1038/s41467-022-28090-5. URL: http://dx.doi.org/10.1038/s41467-022-28090-5.

[53] Adriano B.L. Tort et al. “Temporal Relations between Cortical Network Oscillations and Breathing Frequency during REM Sleep”. In: The Journal of Neuroscience 41.24 (May 2021), pp. 5229–5242. ISSN: 1529-2401. DOI: 10.1523/jneurosci.3067-20.2021. URL: http://dx.doi.org/10.1523/JNEUROSCI.3067-20.2021.

[54] A. Mouraux and G.D. Iannetti. “Across-trial averaging of event-related EEG responses and beyond”. In: Magnetic Resonance Imaging 26.7 (Sept. 2008), pp. 1041–1054. ISSN: 0730-725X. DOI: 10.1016/j.mri.2008.01.011. URL: http://dx.doi.org/10.1016/j.mri.2008.01.011.

[55] Gregory Lee et al. “PyWavelets: A Python package for wavelet analysis”. In: Journal of Open Source Software 4.36 (Apr. 2019), p. 1237. ISSN: 2475-9066. DOI: 10.21105/joss.01237. URL: http://dx.doi.org/10.21105/joss.01237.

[56] The Good Scents Company. The Good Scents Company Information System. http://www.thegoodscentscompany.com. Accessed: 2026-01-15. 2026.

[57] Catharine H. Rankin et al. “Habituation revisited: An updated and revised description of the behavioral characteristics of habituation”. In: Neurobiology of Learning and Memory 92.2 (Sept. 2009), pp. 135–138. ISSN: 1074-7427. DOI: 10.1016/j.nlm.2008.09.012. URL: http://dx.doi.org/10.1016/j.nlm.2008.09.012.

[58] L Wang. “The correlation between physiological and psychological responses to odour stimulation in human subjects”. In: Clinical Neurophysiology 113.4 (Apr. 2002), pp. 542–551. ISSN: 1388-2457. DOI: 10.1016/s1388-2457(02)00029-9. URL: http://dx.doi.org/10.1016/S1388-2457(02)00029-9.

[59] Kwangsu Kim et al. “Odor habituation can modulate very early olfactory event-related potential”. In: Scientific Reports 10.1 (Oct. 2020). ISSN: 2045-2322. DOI: 10.1038/s41598-020-75263-7. URL: http://dx.doi.org/10.1038/s41598-020-75263-7.

[60] Maria Paola Cecchini et al. “Olfaction in patients with Parkinson’s disease: a new threshold test analysis through turning points trajectories”. In: Journal of Neural Transmission 128.11 (July 2021), pp. 1641–1653. ISSN: 1435-1463. DOI: 10.1007/s00702-021-02387-z. URL: http://dx.doi.org/10.1007/s00702-021-02387-z.

[61] Hirokazu Kumazaki et al. “Brief Report: Olfactory Adaptation in Children with Autism Spectrum Disorders”. In: Journal of Autism and Developmental Disorders 49.8 (May 2019), pp. 3462–3469. ISSN: 1573-3432. DOI: 10.1007/s10803-019-04053-6. URL: http://dx.doi.org/10.1007/s10803-019-04053-6.

[62] J. McLinden et al. “Phase-Amplitude Coupling Between EEG Cortical Oscillations and Respiration: An Exploratory Study”. In: 2023 11th International IEEE/EMBS Conference on Neural Engineering (NER). IEEE, Apr. 2023, pp. 1–4. DOI: 10.1109/ner52421.2023.10123888. URL: http://dx.doi.org/10.1109/NER52421.2023.10123888.

[63] John J. Foxe and Adam C. Snyder. “The Role of Alpha-Band Brain Oscillations as a Sensory Suppression Mechanism during Selective Attention”. In: Frontiers in Psychology 2 (2011). ISSN: 1664-1078. DOI: 10.3389/fpsyg.2011.00154. URL: http://dx.doi.org/10.3389/fpsyg.2011.00154.

[64] Claire Martin et al. “Learning-induced modulation of oscillatory activities in the mammalian olfactory system: The role of the centrifugal fibres”. In: Journal of Physiology-Paris 98. 4-6 (2004), pp. 467–478. ISSN: 0928-4257. DOI: 10.1016/j.jphysparis.2005.09.003. URL: http://dx.doi.org/10.1016/j.jphysparis.2005.09.003.

