## Supplemental Material for "Respiratory Phase Orchestrates Human Olfactory Cortical Dynamics"

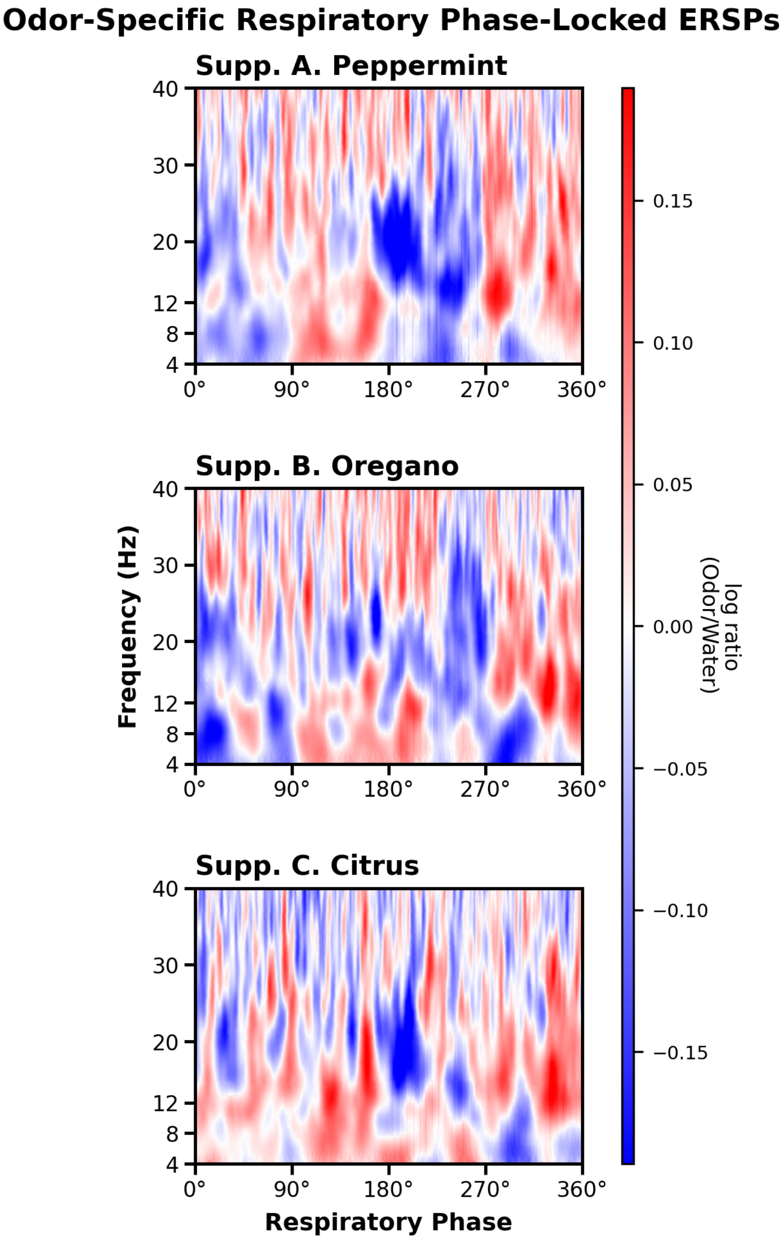


Figure S1: **Odor-specific grand-average respiratory phase-locked event-related spectral perturbations (RPL-ERSPs).**
Grand-average RPL-ERSPs across all subjects ($n = 17$) plotted across the full respiratory cycle ($0^\circ$ = inhalation onset; $180^\circ$ = exhalation onset) separated by stimulus condition. Panels display the phase-locked spectral structure for **Supp. A:** Peppermint, **Supp. B:** Oregano, and **Supp. C:** Citrus. Across conditions, distinct phase-dependent dynamics—characterized by theta-band increases during inhalation and alpha-band decreases during exhalation—persist at the group level when accounting for individual variations in respiratory timing across odor stimuli.

### 1 Supplemental

#### 1.1 Design Choices

To dispense scented air to the subject, a commercially available air compressor, the Spriek 'Intelligent Digital Display Nebulizer for Adults and Kids', was used. This choice was based on its affordability, low noise level ($\leq 50\,\mathrm{dB}$), adequate air delivery, and user-friendly features. Various air compressor nebulizers may be suitable, provided they have universal connection tubing and sufficient air supply. However, it is crucial to avoid excessive air flow, as concentrations > $5\,\mathrm{L/min}$ can cause nasal drying and irritation without proper temperature or humidity control Lötsch et al. 1998. For nasal cannula delivery without these controls, a maximum flow rate of approximately $3\,\mathrm{L/min}$ is recommended to prevent discomfort Lundström et al. 2010.

As a proof of concept, three odors and one no-odor control were chosen for this olfactometer. Air flow branches were linked using "Y" connectors in a binary tree structure. Each branch was connected to a flowmeter Fig.**??C**, allowing individual or uniform flow between $0$ to $5\,\mathrm{L/min}$. Consistent airflow across scent containers ensures constant odor delivery, while flow rates determine the quantity of odorants delivered.

After optimizing flow rates, air was directed through a bubble humidifier (Fig.**??D**) to add water vapor to the airstream without requiring heating. When passing high flow rates through bubble humidifiers, they are unable to create the necessary amount of humidity, due to decreased contact times. In contrast, when passing lower flow rates of <$10\,\mathrm{L/min}$, the gas effectively passes through the water from the bottom of the liquid to the surface, and this contact results in increased water vapor Dasgupta, Ghosh, and Chandra 2022. Thus, a lower flow rate of $0$ to $5\,\mathrm{L/min}$ was used.

The bubble humidifiers (odor containers) were constructed from common Ball® wide-mouth mason jars, due to their glass composition (a nonporous material), affordability, and ease of cleaning. The glass jars were fitted with Tygon tubing, 7.94mm ID x 12.7mm OD for the outlet, and 9.53mm ID x 15.88mm OD for the inlet. To mitigate danger of injury to the patient and the experimenter, safety pop-off pressure valves rated for $1 - 5$ PSI were integrated into the design. Precise holes were drilled for the tubing and the valve using a metal laser cutter. For this design, temperature control of the air was not integrated. Although heated humidifiers increase moisture, there is a trade-off with increased heat allowing for more bacteria colony clustering Koenig and Truwit 2006; Al Ashry and Modrykamien 2014. One odor container served as a control with DI water (no odor). Others contained a precise quantity of essential oil diluted in DI water. Glass bubble humidifiers were chosen for patient comfort, simplicity, cost-effectiveness, and hygiene. The housing for the odor containers was 3D printed. The design allows for multiple odors containers in groups of 2, and space between the top and bottom holder for flowmeter and tubing placement.

After the humidifier, scented air or control air passes through a 3-way solenoid valve, which directs residual scented air outside of the experimentation room when not in use Fig.**??E**, eliminating the need for expensive air reservoirs. This design prevents pressure build-up and backflow. A 3-way solenoid valve operates by having ports 1 & 2 usually connected, when the solenoid is not powered, and switching to connect ports 1 & 3 when the solenoid is powered. In this way, our design allows scented air to pass through tubes to outside the experimentation room when an odor is not intended for presentation, while allowing scented air to travel towards the patient when the solenoid is powered.

After the first solenoids where the desired odor was selected, the four scent lines merge. Immediately before the tube to the nasal cannula, another 3-way solenoid is connected, allowing for precise control of timing before reaching the nasal cannula. Excess air is again directed outside the room.

Nasal cannulas were chosen as the ideal delivery device for this experiment. They offer precise control over odor timing and concentration, are cost-effective, disposable, and can be purchased with nonstick materials like Teflon B. Johnson, Khan, and Sobel 2008. Nasal cannulas allow directed airflow to both nostrils without relying on active sniffing, which has been shown to improve olfactory stimulus localization Al Aïn and Frasnelli 2017. While other methods like scented bottles, papers, or fan systems were considered Lorig and Schwartz 1988; Martin 1998; Krbot Skorić et al. 2014; Sowndhararajan et al. 2016; Kim et al. 2019, they introduce temporal issues such an imprecise onset/offset timing, along with inability to control concentration or distance from scent effectively.

##### 1.2 Electronics

The precision of this olfactometer, in delivering measured and controlled volumes of scented air for a defined duration, is attributed to its integrated electronics. A diode was connected in parallel to the solenoid controlling airflow direction to protect the switching transistor. Upon deactivation, solenoids attempt to maintain current flow; these diodes provide a unidirectional path for this current, thereby protecting the rest of the circuit from potential damage. To mitigate 60Hz noise challenges with the sensitive EEG system, the solenoids are powered by an external 2500mAh lithium-ion battery.

An Arduino control module enhances versatility by permitting easy integration with external devices, including the EEG system. For instance, Transistor-Transistor-Logic (TTL) pulses can be transmitted via Arduino for precise timing synchronization between odor delivery and neural activity. TTL pulses, typically represented as digital '0' (LOW) or '1' (HIGH) states Artoni et al. 2018, enable external triggering. This allows for applications such as marking odor release events directly aligned to the EEG data, resulting in straightforward post-experiment analysis of relevant neural epochs.

The Arduino Uno's *digitalRead* and *digitalWrite* commands operate with an average latency of approximately 3.4 µs RoboticsBackEnd 2021, ensuring that TTL pulses are sent with minimal delay for accurate time stamping.

##### 1.3 Nasal Respiration Sensor

Accurate millisecond-level odor presentation in olfactometry critically depends on real-time respiratory feedback. Olfaction is fundamentally a respiratory-dependent process, as inhales are required to transport odorants to the olfactory epithelium. Therefore, even a precisely timed odor delivery may go unnoticed or be substantially delayed if it coincides with exhalation. While past studies have utilized methods like thoracic or abdominal respiratory belts to monitor respiration, research indicates that sensors placed proximally to the nose provide more precise measurements of airflow dynamics B. N. Johnson et al. 2006; Zelano et al. 2016. For accurate temperature and pressure sensor readings, a small flexible PCB was fabricated incorporating a Bosch BME280 combination temperature/pressure/humidity sensor, which was inserted into the center of a nasal cannula, between the two nostrils. Fig.**??** demonstrate functionality of the normalized humidity recordings for one subject and one trial. The peaks indicate the start of inhalation, and the troughs indicate the onset of exhalation. The bold curve is the averaged curve across trials for one subject. This figure demonstrates that the TTL pulse was triggered on the onset of inhale, insuring the optimal release of scented air to the subject at the desired point in time.

The precise odor delivery protocol, synchronized with individual respiration, involves the following four key stages:

1. **Breath Detection**: The Arduino monitors breathing by measuring temperature and humidity changes. Inhales are detected by decreasing values, exhales by increasing values.
2. **Odor Preparation**: Following a random inter-stimulus interval of 15-30 seconds, the first solenoid activates for up to 10 seconds, drawing and preparing a specific odorant for delivery.
3. **Scent Delivery Timing and Verification**: Upon the subject's first detected exhalation immediately after an inhalation, the second solenoid opens for one second. This precise timing allows the prepared scented air to traverse the 1.21 meter nasal cannula and reach the subject's nose at the onset of their subsequent inhalation.
4. **EEG Synchronization and Scent Arrival Timestamp**: A TTL pulse is transmitted to the EEG system, recording a precise timestamp at the moment the subject begins their subsequent detected inhalation. This pulse marks the exact point when the scented air could enter the nasal cavity, accounting for minimal electrical and mechanical delays (on the order of microseconds).

This design ensures that the scent reaches the subject at the optimal moment in their breathing cycle, allowing for precise control and measurement in olfactory experiments.
